# Differential seasonal effects of ephemeral legumes in response to moisture-nitrogen coupling in the deserts of northwestern China

**DOI:** 10.1101/2024.03.19.585727

**Authors:** Yuxin Xiao, Boyi Song, Jinqiu Li, Nargiza Galip, Ao Yang, Xinyu Zhang, Weiwei Zhuang

## Abstract

Desert ecosystems are ecosystems limited by both nitrogen and moisture, while legumes, as an important source of nitrogen in desert ecosystems, are extremely sensitive to respond to moisture and nitrogen changes. In order to reveal the growth and physiological metabolic processes of desert legumes in response to moisture and nitrogen changes. In this study, two dominant ephemeral legumes, *Trigonella arcuata* C. A. Mey. and *Astragalus arpilobus* Kar. & Kir. were selected from the Gurbantunggut Desert through a controlled experiment in greenhouse pots: four moisture treatment levels were established: W1(soil moisture content of 5 %), W2(soil moisture content of 7 %), W3( 9 % soil moisture content), W4(11 % soil moisture content) and three N treatment levels: N1(0 mmol/L ^15^NH_4_ ^15^NO_3_), N2(18 mmol/L ^15^NH_4_ ^15^NO_3_), N3(72 mmol/L ^15^NH_4_ ^15^NO_3_), and we conducted a ^15^N tracking experiment for the pot treatments throughout the growing season. The results showed that: (1) moisture and nitrogen treatments had similar patterns on above-ground and below-ground as well as total fresh weight, plant height and basal diameter, Chla, Chlb and Chla+b of the two legumes, all of which were greatest in the W3N3 treatment and smallest in the W4N1 treatment, but the growth characteristics and chlorophyll parameters had cumulative differences over the growing season. (2) When subjected to severe moisture and nitrogen stresses, the 2 plants would increase the physiological levels of stress tolerance in vivo to enhance cellular resistance, with the osmoregulatory substance content being the most sensitive in response to moisture, and the cell membrane and antioxidant enzyme systems being the most sensitive in response to nitrogen stress. (3) The adaptive characteristics of physiological metabolic processes in plants to seasonal changes, in which growth characteristics, cell membrane and antioxidant enzyme content accumulation showed: rapid growth period < maximum biomass period, while chlorophyll content and osmoregulatory substance content accumulation content accumulation showed: rapid growth period > maximum biomass period. In general, drought can hinder nitrogen uptake and transport, while sufficient moisture can accelerate the nutrient metabolism process of plants, and there is a nutrient dilution effect once the moisture is in excess, but in the nitrogen starvation state, the moisture factor has almost no effect on plants, therefore, the moisture-nitrogen coupling effect is more reflective of the physiological response characteristics of the plants when they encountered the adversity stress, and the degree of accumulation is different in different growing seasons, and the plants in the The differences in different growing seasons reflect the changes in photosynthetic intensity and energy demand.

## Introduction

In the context of global climate change, climate change-induced ecosystem vulnerability is increasing and causing profound implications on ecosystem structure and functioning. The Intergovernmental Panel on Climate Change(IPCC) in its Sixth Assessment Report highlights a projected trajectory of rising average surface temperatures and precipitation for the Central Asian region(IPCC, 2021). The CMIP5 Global Climate Model simulations predict a continuation of “warming and humidification” under future climate scenarios, which will have a significant impact on the spatial and temporal patterns of dominant herbaceous plant taxa in the Central Asian desert ecosystems. Notably, Xinjiang, as a pivotal area within the Northern Hemisphere’s “warming and humidification” belt, stands out for its unique position in the mid-latitudes(Shi et al., 2007), and the intra-annual uniformity of precipitation distribution will tend to be homogenised, which affects the desert ecosystems of Central Asia, and has an important impact on the spatial and temporal patterns of the dominant herbaceous plant groups in this ecoregion. In addition to precipitation alterations due to climate change, nitrogen deposition is increasingly becoming a serious threat, e.g., the annual mean nitrogen deposition in agroecosystems in the arid zone of Xinjiang can reach up to 19.5 kg/hm^2^(Li et al., 2013), and that in the oases around the desert can reach as high as 30 kg/hm^2^(Xu et al., 2015; Zhou et al., 2014), exceeding many ecosystem nitrogen thresholds(10 kg/hm^2^). Global climate change and human activities not only affect the natural environment, ecosystem balance, and various ecohydrological processes, but are also key factors for measuring and predicting ecosystem changes, which are relevant to the biogeocirculation processes of almost all types of landscapes and ecosystems on Earth(Daunoras et al., 2024; Higgins et al., 2023; Koltz et al., 2018; Manlick et al., 2023; Zhang et al., 2019).

Desert ecosystems, as one of the most vulnerable ecosystems on earth, are extremely sensitive in responding to climate change(Hooper and Johnson, 1999). In desert ecosystems, moisture availability is a major limiting factor for plant growth and ecosystem productivity(Huang et al., 2016). Precipitation variability and plant moisture availability directly determine seed germination, seedling establishment, individual morphogenesis, adult maturation and reproductive processes in desert plants, further affecting community structure and ecosystem stability. As the most critical limiting factor in arid and semi-arid areas, a moderate increase in precipitation will increase the photosynthetic capacity of plants(Zhou et al., 2011) and the net primary productivity of vegetation(Guo et al., 2017; Huston et al., 2000), while drought will inhibit plant growth, change the ratio of dry matter distribution, and reduce the accumulation of dry matter(Fereres and Moreno, 2008; Lassouane et al., 2013). It has been suggested that drought stress promotes the uptake of nutrients by plants, which in turn promotes the acquisition of moisture by plants to alleviate drought stress(Zhou et al., 2011). It has also been suggested that moisture changes alter the intrinsic nutrient niche acquisition strategies of desert plants(Xiao et al., 2023). Besides moisture, nitrogen is the second most important limiting factor affecting desert ecosystems. The effects of nitrogen addition on biomass are mainly significant increase, no effect and significant decrease(Su et al., 2013; Wu et al., 2016). Compared with other ecosystems such as forests, desert ecosystems plant productivity, community composition and structure respond more significantly to short-term atmospheric nitrogen deposition(Clark and Tilman, 2008; Pernetta, 2012). Some studies have shown that nitrogen deposition can increase the cover of herbaceous plants and decrease the abundance of legumes in the northern region of Chihuahuan Desert, thus changing the structure of plant communities(Baez et al., 2007). In fact, the effects of nitrogen deposition on plant and microbial community activities are usually regulated by moisture(Holt and Chesson, 2014). Therefore, studying the coupled effects of moisture and nitrogen factors or other multifactorial joint effects can more comprehensively explain the relationship between plant growth and changes in environmental factors.

Legumes are widely distributed globally and are the third largest family of angiosperms after Asteraceae and Orchidaceae, and are one of the few plant groups with symbiotic nitrogen fixation. Nitrogen fixation by legumes is an important source of nitrogen in terrestrial ecosystems, accounting for about 80% of global biological nitrogen fixation. Its symbiosis with rhizobia converts atmospheric nitrogen into plant-available nitrogen compounds(Notaris et al., 2019). This process is particularly important for N-poor soils, increasing soil fertility and promoting the growth of other non-nitrogen-fixing plants in the ecosystem(Vitousek et al., 2013; Zhang et al., 2019). Legumes are widely distributed in arid zone ecosystems such as deserts and are the centre of ecosystems providing effective nitrogen inputs(Richard, 1982), occupying an extremely important position in desert ecosystems. In addition to this, the growth and nitrogen fixation of legumes are highly susceptible to environmental factors. It has been pointed out that precipitation changes affect the inter-root microbial changes and nodulation characteristics of legumes, influencing the availability of nitrogen and the rate of ion transport(Kreuzwieser and Gessler, 2010), which in turn affects their nitrogen fixation, and likewise their coexistence and competition with neighbouring plants(Dhamala et al., 2017; Sadras et al., 2016). In addition to moisture, increased exotic nitrogen is increasingly becoming an important factor affecting legumes. It has been shown that despite their nitrogen fixation, legumes are significantly lower than non-nitrogen fixing plants in terms of nitrogen utilisation efficiency and growth rate response to nitrogen deposition(Chesnais et al., 2016; Shinano et al., 2012). Although legumes are able to fix atmospheric nitrogen through rhizobia, the nitrogen level in the soil still affects their growth. For example, soil substrate also affects plant nutrient uptake and rhizobial nitrogen-fixing capacity, and high levels of soil nitrogen typically inhibit rhizoma formation in legumes, as plants tend to take up available nitrogen in the soil directly rather than relying on energy-intensive nitrogen-fixation processes(Vitousek et al., 2013). While legumes have a special contribution to desert ecosystems as an important source of effective nitrogen inputs, their physiological traits and nitrogen fixation characteristics in response to changes in environmental factors are also extremely sensitive, which can in turn affect the ecosystem again.

The Gurbantunggut Desert, located in the Junggar Basin of Xinjiang, China, is the largest fixed and semi-fixed desert in China(Zhou et al., 2014). As a typical pioneering species of nitrogen-fixing plants in this region, the changing characteristics of legumes in response to environmental factors are of great significance in assessing the effects of nitrogen cycling in desert ecosystems in this region. In the Gurbantunggut Desert, ephemeral plant taxa occupy up to 70-80% of the herbaceous plant biomass, which is very important for the stability of sand dunes in the early spring season(Zhou et al., 2014). Ephemeral plants in this desert are more sensitive to moisture, temperature and other environmental factors compared to other living plants. It has been suggested that the selective distribution of ephemeral plants to dune sites in this region is mainly dependent on soil moisture conditions(Xiao et al., 2023). There are 25 species of ephemeral legumes in the Gurbantunggut Desert, which are important in the vegetation cover and nitrogen supply of this desert ecosystem. Based on the importance of ephemeral legumes in desert ecosystems and the threat of altered precipitation and increased nitrogen deposition to desert ecosystems, we ask the scientific question: how do the growth and physiological processes of ephemeral legumes respond to changes in moisture and nitrogen environmental factors in a typical temperate desert ecosystem? Due to the global and long-term nature of precipitation and atmospheric nitrogen deposition, its impact on desert ecosystems is also a long-term and intricate process(Huang et al., 2016), and few similar studies on desert ecosystems have been reported. Therefore, we propose the hypothesis that: (1) Compared with single moisture and nitrogen stresses, coupled moisture and nitrogen effects may alter the original moisture use and nitrogen acquisition strategies of desert ephemeral legumes. (2) When experiencing moisture and nitrogen stress, plants may flexibly regulate their response mechanisms, with different physiological indicators responding to different environmental factors with different degrees of sensitivity. (3) Due to the short life history of ephemeral plants and their extreme sensitivity to environmental changes, the physiological and metabolic processes of ephemeral legumes may differ in response to moisture and nitrogen stresses in different growing seasons. Accordingly, we chose two typical dominant species in this desert, the ephemeral plants *Trigonella arcuata* and *Astragalus arpilobus*, as our research subjects, and through indoor potting experiments simulating coupled moisture and nitrogen stress, we attempted to comprehensively analyse the coupled effects of moisture and nitrogen on the growth of ephemeral legumes in deserts, from phenological characteristics to physiological and biochemical levels, and to assess the seasonal contribution of nitrogen in legumes. This study will provide a theoretical basis for understanding the response of desert legumes to nitrogen deposition and precipitation under global climate and environmental changes, and the nitrogen cycle in the region.

## Material and methods

### Experimental materials

In June 2020, seeds from two legume species were collected from their natural habitats, amounting to 400-500 seeds per species. Subsequently, these seeds underwent a vernalization process in the laboratory for 30 days in March of the following year. The growth substrate was also selected from desert habitat soil (soil organic carbon, total nitrogen, total phosphorus, N-NH_4_^+^, N-NO_3_^-^ and pH were 1.72 g·kg^-1^, 0.18 g·kg^-1^, 0.51 g·kg^-1^, 14.28 μg·g^-1^, 22.05 μg·g^-1^, 7.01), apoplastic and larger particles were removed using a 2 mm funnel sieve, and then mixed thoroughly and evenly. The test soil was mixed and packed into pots (height 25 cm, upper diameter 28 cm), each containing 7.0 kg of soil, and 20 seeds were sown in each pot. Seedlings were thinned out after 2 weeks of germination, and only four well-developed seedlings with little variation in shape were retained in each pot as test material. The pot culture was carried out in a greenhouse and all pots were randomly placed in the greenhouse.

### Experiment design

Based on the meteorological data and the current status of nitrogen deposition in the Junggar Desert in northern Xinjiang in the last recent years, four moisture treatment levels were established: W1(soil moisture content of 5%), W2(soil moisture content of 7%), W3(soil moisture content of 9%), and W4(soil moisture content of 11%), and three nitrogen treatment levels: N1(0 mmol/L ^15^NH_4_ ^15^NO_3_), N2(18 mmol/L ^15^NH_4_ ^15^NO_3_), and N3(72 mmol/L ^15^NH_4_ ^15^NO_3_), where moisture treatments were applied once every 3 d using the weighing method, nitrogen additions were applied once every 1 week using the more advanced ^15^N isotope-tracking technique, and the total amount of N_2_ treatments was the critical nitrogen concentration affecting the majority of the ecosystems(about 1 g N · m^-2^ · a^-1^), and the N3 treatments were The total amount of applied nitrogen was equivalent to the nitrogen deposition near the Gurbantunggut Desert(about 0.8 g N · m^-2^ · a^-1^)(Li et al., 2013) and the maximum nitrogen deposition in the Mojave Desert, USA(about 3.23 g N · m^-2^ · a^-1^)(Brooks, 2003), respectively. Plants were harvested whole at the rapid growth period(45th day after the emergence of the second true leaf) and at the maximum biomass period(90th day after the emergence of the second true leaf) of the two species, respectively.

### Determination of chlorophyll fluorescence parameters

Junior-Pam portable modulated chlorophyll fluorescence meter was used to measure chlorophyll fluorescence parameters. The actual photochemical efficiency YII and the parameter data needed to calculate the maximum photochemical efficiency Fv/Fm were obtained under normal light conditions and after dark adaptation for 30 min respectively(under normal light adaptation: YII=(Fm’-Fs)/Fm’, after dark adaptation for 30 min: Fv/Fm=(Fm-Fo)/Fm)

### Determination of growth characteristics

Before harvesting, plant height was measured using a tape measure and basal stem was measured using a vernier caliper(3 plants were measured and averaged). Four plants were harvested from each pot, the whole plant was dug up and the roots were rinsed with distilled moisture to remove the sandy soil and blotted with filter paper, and the roots, stems and leaves were separated with scissors. The fresh weight of each sample was weighed using a 10,000 parts per million balance, and the combined weight of the fresh stems and fresh leaves of the same sample was recorded as above ground fresh weight(AGB), the weight of the fresh roots was recorded as below ground fresh weight(BGB), the sum of the three was recorded as total fresh weight(TB), and the ratio of the below ground fresh weight(BGB) to the above ground fresh weight(AGB) was recorded as the root/crown ratio(R/S).

### Determination of chlorophyll content

Chlorophyll content (Chl) was determined using anhydrous ethanol-acetone(2: 1 v/v) mixture leaching method.

### Determination of osmoregulatory substances

Free proline content (Pro) was determined using the acid ninhydrin method, soluble sugar content (SS) using the anthrone colourimetric method and soluble protein content (SP) using the Caumas Brilliant Blue method(Lassouane et al., 2013; Monreal etal., 2007).

### Determination of cell membrane and antioxidant enzyme system

The content of Malondialdehyde (MDA) was assessed employing the Thiobarbituric Acid (TBA) assay, while the activity of Superoxide Dismutase (SOD) was measured through the hydroxylamine method. Furthermore, the activity of Peroxidase (POD) in moss was determined utilizing the guaiacol method, and Catalase (CAT) activity was quantified via the ammonium molybdate technique(Sun et al., 2009; Wu et al., 2012).

### Statistical analysis

We first checked the normality of the data using SPSS20.0(Chicago, IL, USA), The general linear model and Duncan analysis method were used to analyze AGB, BGB, TB, R/S, Chla, Chlb, Chla+b, Chla/b, YII, Fv/Fm, Pro, SS, SP, MDA, SOD, POD and CAT of the two species by one-way ANOVA and comparison of difference significance. Four-factor analysis of variance was used to analyze the effects of variation sources(species, growing season, moisture, nitrogen and their interactions) on the above indicators, and the difference significance level was 0.05. Path analysis was conducted using partial least squares path modelling(PLS-PM). The calculation of the model was performed using the package ‘plspm’ in R version 3.5.2(ver. 0.4.9, Sanchez, 2013). The correlation matrix between variables was analysed using principal component analysis(PCA). All figures were drawn on Origin 2021.

## Results

### Growth and biomass allocation characteristics

Considering four sources of variation: species(S), growing season(G), moisture(M), nitrogen(N), and their interactions, this study analyzes changes in various growth characteristics such as aboveground, belowground and total fresh weight, root-shoot ratio, plant height, and basal diameter of two leguminous plants. As shown in Table 1, each factor has varying degrees of impact on the growth indices including aboveground, belowground and total fresh weight, root-shoot ratio, plant height, and basal diameter for both *Trigonella arcuata* and *Astragalus arpilobus*. Among these factors, moisture and nitrogen have a highly significant effect(*P* < 0.001) on all growth indices.

**Table 1.**
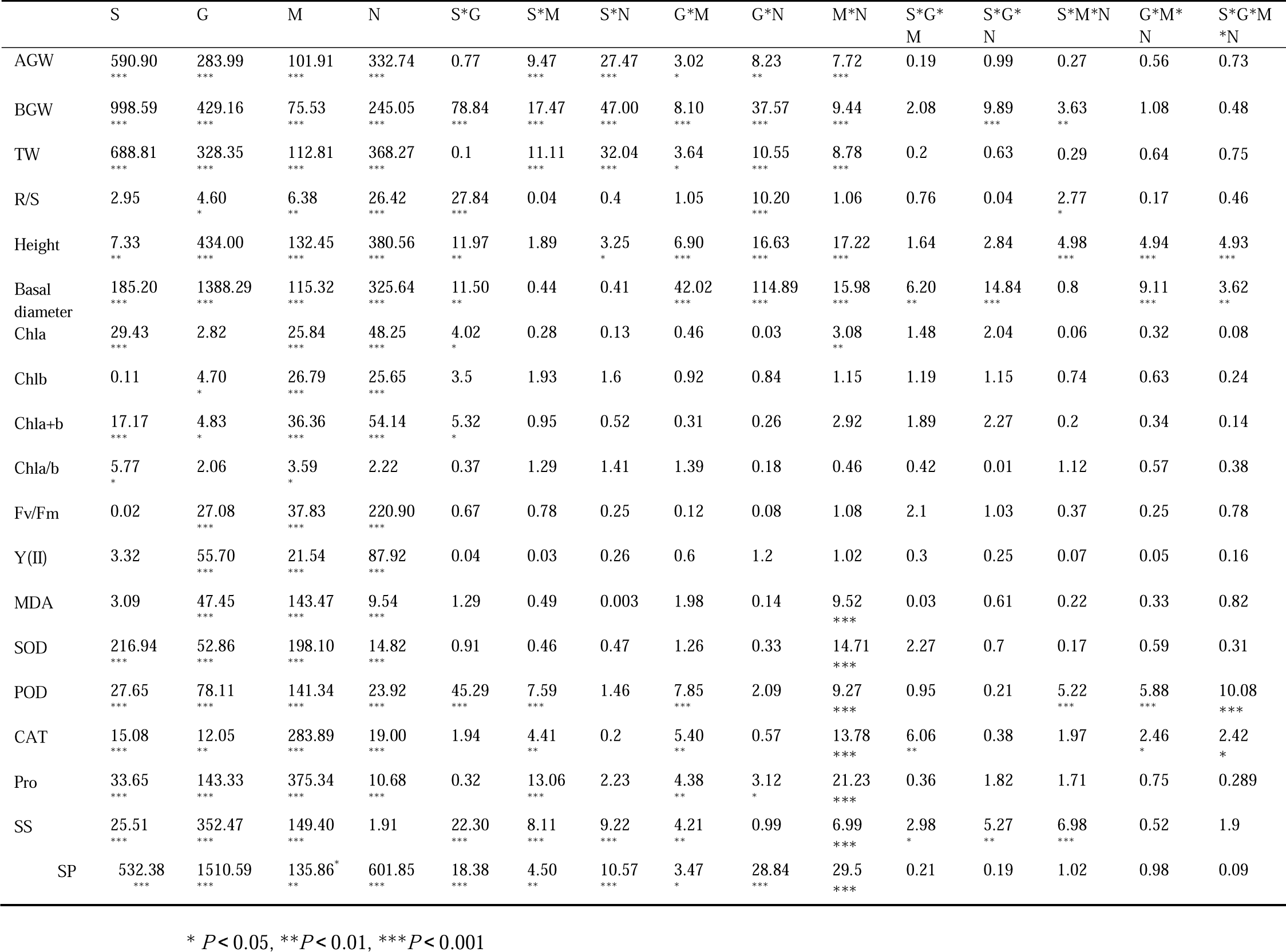
Four-factor ANOVA of the effects of species, growing season, moisture, nitrogen, and their interactions on the growth and physiological indicators of two desert leguminous plants.

Through the analysis of the effects of various factors on the fresh weight and root-shoot ratio of two plants, it was found that species, growth season, moisture, nitrogen, species × moisture, species × nitrogen, growth season × moisture, growth season × nitrogen, and moisture × nitrogen significantly influenced(*P* < 0.001, *P* < 0.01, *P* < 0.05) the aboveground, belowground, and total fresh weight of *Trigonella arcuata* and *Astragalus arpilobus*. The belowground fresh weight was more significantly affected by the interactions between factors, with species × growth season, growth season × moisture, growth season × nitrogen, species × growth season × nitrogen, and species × moisture × nitrogen showing highly significant effects(*P* < 0.001, *P* < 0.01) on the belowground fresh weight of both plants. Furthermore, the root-shoot ratio was primarily influenced by the growth season, moisture, nitrogen, species × growth season, growth season × nitrogen, and species × moisture × nitrogen (*P* < 0.001, *P* < 0.01, *P* < 0.05). Further analysis of the key factors, moisture and nitrogen, on the fresh weight and root-shoot ratio of both plants revealed(see Fig. 1) that under the same nitrogen addition treatment, the aboveground, belowground, and total fresh weight of both plants were affected by different moisture gradients, showing the trend: W3 > W2 > W4 > W1. Conversely, under the same moisture levels, the influence of different nitrogen levels on the aboveground, belowground, and total fresh weight of both plants was: N1 < N2 < N3. Combining the effects of moisture-nitrogen interactions revealed that the accumulation of all growth indicators was manifested as follows: rapid growth period < maximum biomass period. The maximum aboveground, belowground, and total fresh weight were recorded in the W3N3 treatment, and the minimum in the W1N1 treatment. During the rapid growth period, the total fresh weight variation was more pronounced across different moisture and nitrogen levels, with the total fresh weight of *Trigonella arcuata* in the W3N3 treatment being 5.4431 times that in the W1N1 treatment, and that of *Astragalus arpilobus* being 5.6008 times. However, compared to the rapid growth period, the fresh weight of both plants was more significantly accumulated during the maximum biomass period, with the total fresh weight per plant residing between 1.5056 and 7.8100 g for *Trigonella arcuata*, and 1.2018 and 4.7446 for *Astragalus arpilobus*. The root-shoot ratios of both plants, influenced by the interaction of moisture and nitrogen, exhibited trends opposite to those of fresh weight changes. Under the same nitrogen treatments, the root-shoot ratios of *Trigonella arcuata* and *Astragalus arpilobus* were mostly influenced by different moisture gradients as follows: W1 > W4 > W3, W2, and under the same moisture gradients, the root-shoot ratios were affected by different nitrogen levels, typically showing: N1 > N2, N3. Considering the combined effect of moisture and nitrogen interaction, it was found that, during both the rapid growth and maximum biomass period, the root-shoot ratio under the N1W1 treatment was significantly higher than in other moisture and nitrogen treatment groups. Additionally, the variation in root-shoot ratio were as follows: rapid growth period > maximum biomass period.

**Fig. 1.**
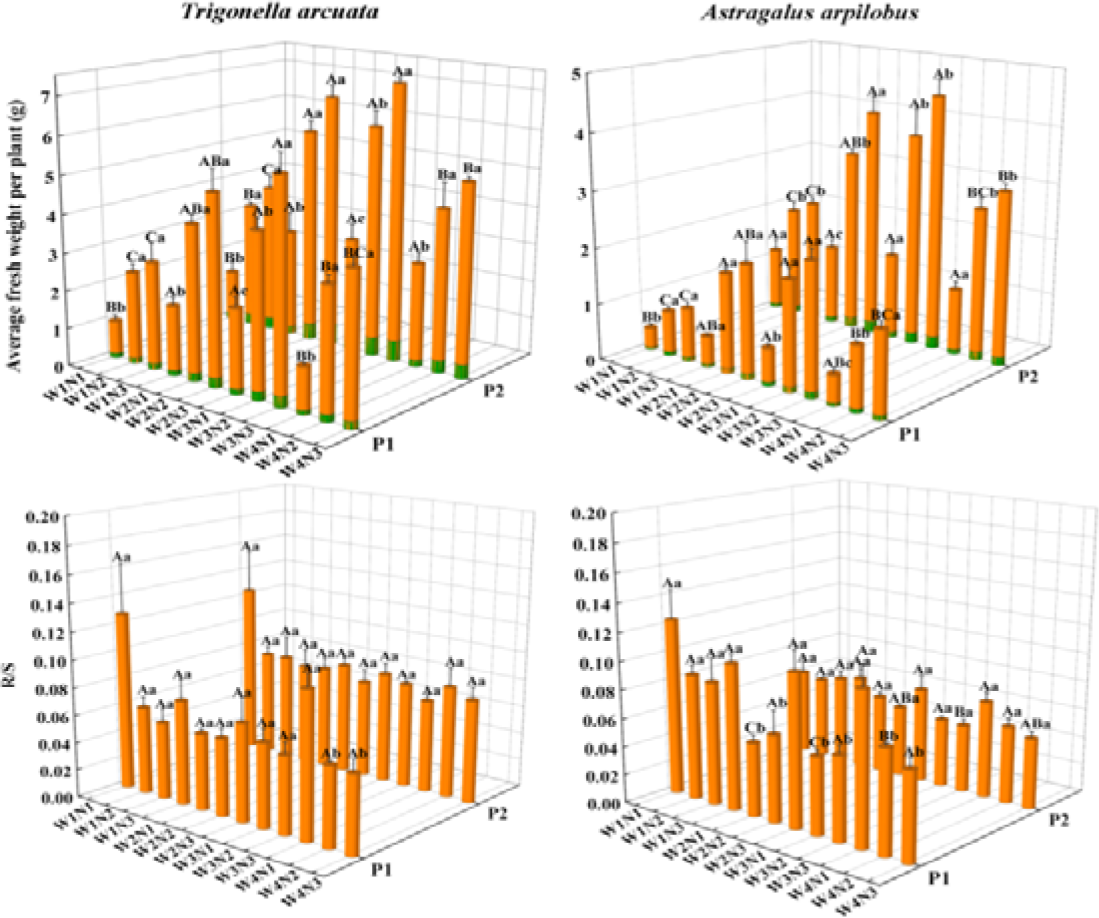
Average above-ground, below-ground, and total fresh weight, and root-shoot ratio of two leguminous plants under the interaction of moisture and nitrogen. P1 represents the rapid growth phase, while P2 denotes the period of maximum biomass. Different uppercase letters indicate significant differences in the same plant under the same nitrogen level across different moisture gradients, while different lowercase letters signify significant differences in the same plant under the same moisture gradient across different nitrogen gradients.

Analysis of the effects of various factors on the plant height and basal diameter characteristics of the two plant species revealed that species, growth season, moisture, nitrogen, species × growth season, growth season × moisture, growth season × nitrogen, moisture × nitrogen, growth season × moisture × nitrogen, and species × growth season × moisture × nitrogen all had a highly significant impact(*P* < 0.001, *P* < 0.01) on the plant height and basal diameter of both plants. Additionally, species × nitrogen and species × moisture × nitrogen had a significant effect(*P* < 0.001, *P* < 0.05) on plant height, whereas species × growth season × moisture and species × growth season × nitrogen significantly influenced(*P* < 0.001, *P* < 0.01) basal diameter. Further analysis of the key factors, moisture and nitrogen, on the fresh weight and root-shoot ratio of the two plants revealed(Fig. 1) that under the same nitrogen addition treatment, the plant height and basal diameter of both species were influenced by different moisture gradients, showing the trend: W3 > W2 > W4 > W1. Similarly, under the same moisture levels, the plant height and basal diameter were affected by different nitrogen levels, typically following the pattern: N1 < N2 < N3, which is consistent with the trend observed in fresh weight changes due to moisture and nitrogen. Combining the effects of water and nitrogen interactions revealed that the accumulation of both growth metrics was expressed as: rapid growth period < maximum biomass period, with the maximum values recorded in the W3N3 treatment and the minimum in the W4N1 treatment(Fig. 2).

**Fig. 2.**
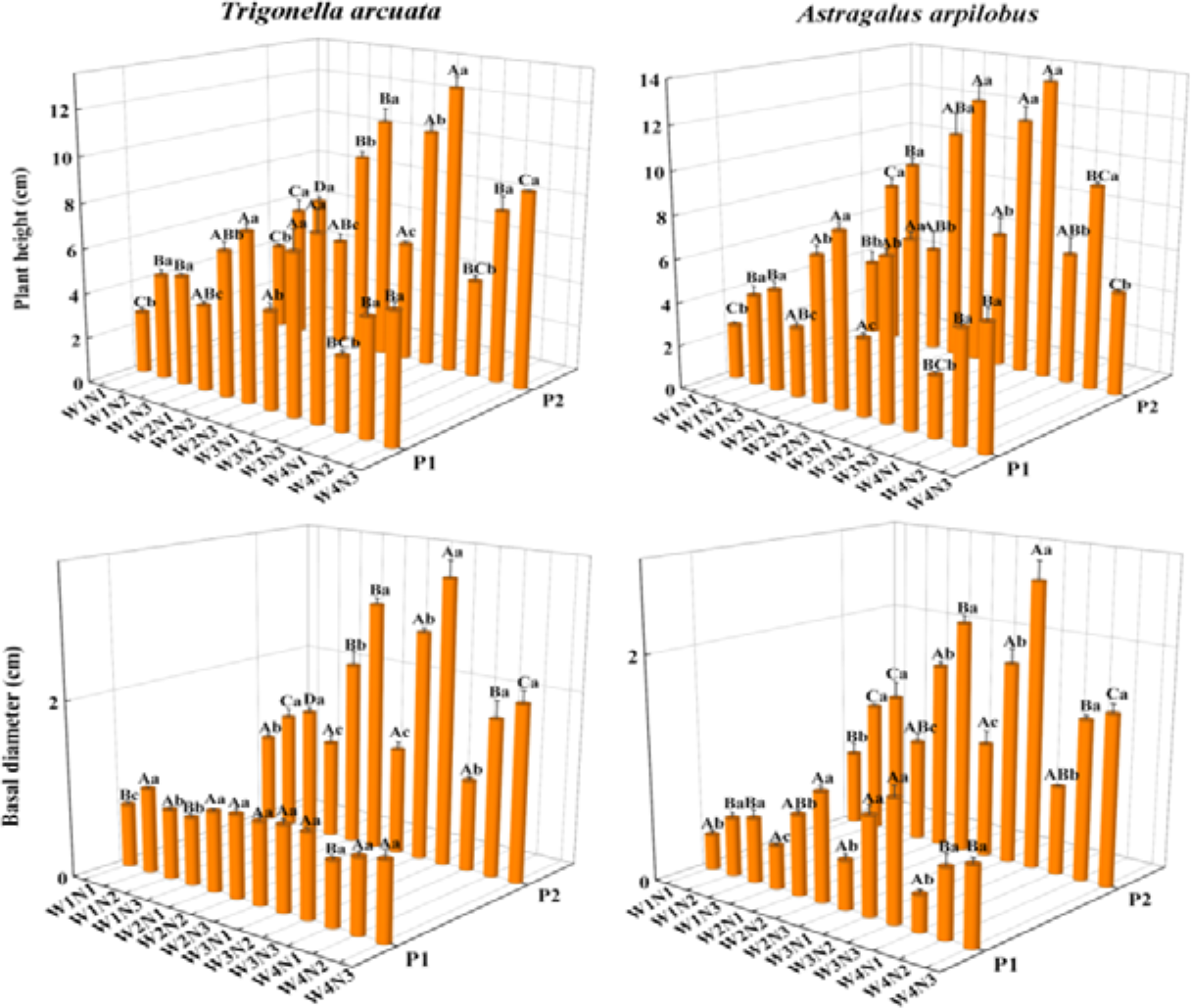
Plant height and basal diameter of two leguminous plants under the interaction of moisture and nitrogen. P1 represents the rapid growth phase, and P2 denotes the period of maximum biomass. Different uppercase letters indicate significant differences in the same plant under the same nitrogen level across various moisture gradients, whereas different lowercase letters denote significant differences in the same plant under the same moisture gradient across different nitrogen gradients.

The results indicate that moisture and nitrogen treatments have similar patterns of influence on the aboveground, belowground, and total fresh weight as well as plant height and basal diameter of the two leguminous plants. Both parameters increased with rising moisture gradients and nitrogen concentrations. However, significant inhibitory effects were observed under dry conditions(W1) and very wet conditions(W4).

### Chlorophyll Fluorescence and Physiological Response

In addition to analyzing the apparent morphological characteristics of potted leguminous plants under the interaction of moisture and nitrogen, we further explored, using physiological and biochemical methods, the response characteristics of the photosynthetic system, osmoregulation system, cell membrane system, and antioxidant enzyme system of potted leguminous plants to changes in moisture and nitrogen. The chlorophyll content of the plant cell photosynthetic system is a key indicator reflecting the plant’ s drought resistance. As shown in Table 1, the species(S), growth season(G), moisture(M), nitrogen(N), and their interactions had consistent patterns of influence on chlorophyll a(Chla), chlorophyll b(Chlb), total chlorophyll(Chla+b), chlorophyll a/b ratio(Chla/b), maximum photochemical efficiency(Fv/Fm), and actual photochemical efficiency(YII) of the two leguminous plants. Species, moisture, nitrogen, species × growth season, and moisture × nitrogen significantly affected Chla(*P* < 0.001, *P* < 0.01, *P* < 0.05). Growth season, moisture, and nitrogen had a significant impact on Chlb(*P* < 0.001, *P* < 0.05), while species, growth season, moisture, nitrogen, species × growth season, and nitrogen significantly influenced Chla+b(*P* < 0.001, *P* < 0.05). Species and moisture significantly affected the Chla/b ratio(*P* < 0.05). Growth season, moisture, and nitrogen had a highly significant effect on Fv/Fm and YII(*P* < 0.001).

Further analysis of the key factors, moisture and nitrogen, on the photosynthetic system of the two plants revealed that under the same nitrogen addition treatment, the Chla, Chlb, and Chla+b of both species were mostly influenced by different moisture gradients, generally showing the trend: W3 > W2 > W4 > W1. However, the Chla/b ratio showed different trends under moisture influence, with *Trigonella arcuata* showing a trend of W1, W4 > W2 > W3 during the rapid growth period, while no consistent pattern was observed in other cases. Under the same moisture treatment, Chla, Chlb, and Chla+b of both plants were generally influenced by different nitrogen levels, showing the trend: N1 < N2 < N3. The Chla/b ratio influenced by nitrogen was similar to moisture, with only *Trigonella arcuata* showing a negative correlation with nitrogen gradient, i.e., the Chla/b ratio decreased as nitrogen gradient increased. *Astragalus arpilobus*’s Chla/b ratio showed no clear trend under the influence of moisture and nitrogen.

Combining the effects of moisture and nitrogen interaction, it was found that the Chla, Chlb, and Chla+b contents of both leguminous plants during the two growth seasons were highest in the W3N3 treatment, and both showed that the rapid growth period > maximum biomass period. During the rapid growth period, the highest Chla, Chlb, and Chla+b contents in *Trigonella arcuata* reached 1.0774, 0.4698, 1.5471 mg/g FW respectively, while in *Astragalus arpilobus*, they reached 0.8648, 0.4207, 1.2855 mg/g FW. In the W3N3 treatment, these values were significantly higher than in the W3N2 treatment, suggesting that adequate nitrogen improves the effective utilization of moisture in leguminous plants. The lowest contents of Chla, Chlb, and Chla+b in both plants were observed in the W1N1 treatment, with the stress effect being more pronounced during the maximum biomass period. In most cases, the Chla/b ratio in both plants was higher in the W1N1 treatment, which is opposite to the trend observed for Chla, Chlb, and Chla+b under the influence of moisture and nitrogen(Fig. 3).

**Fig. 3.**
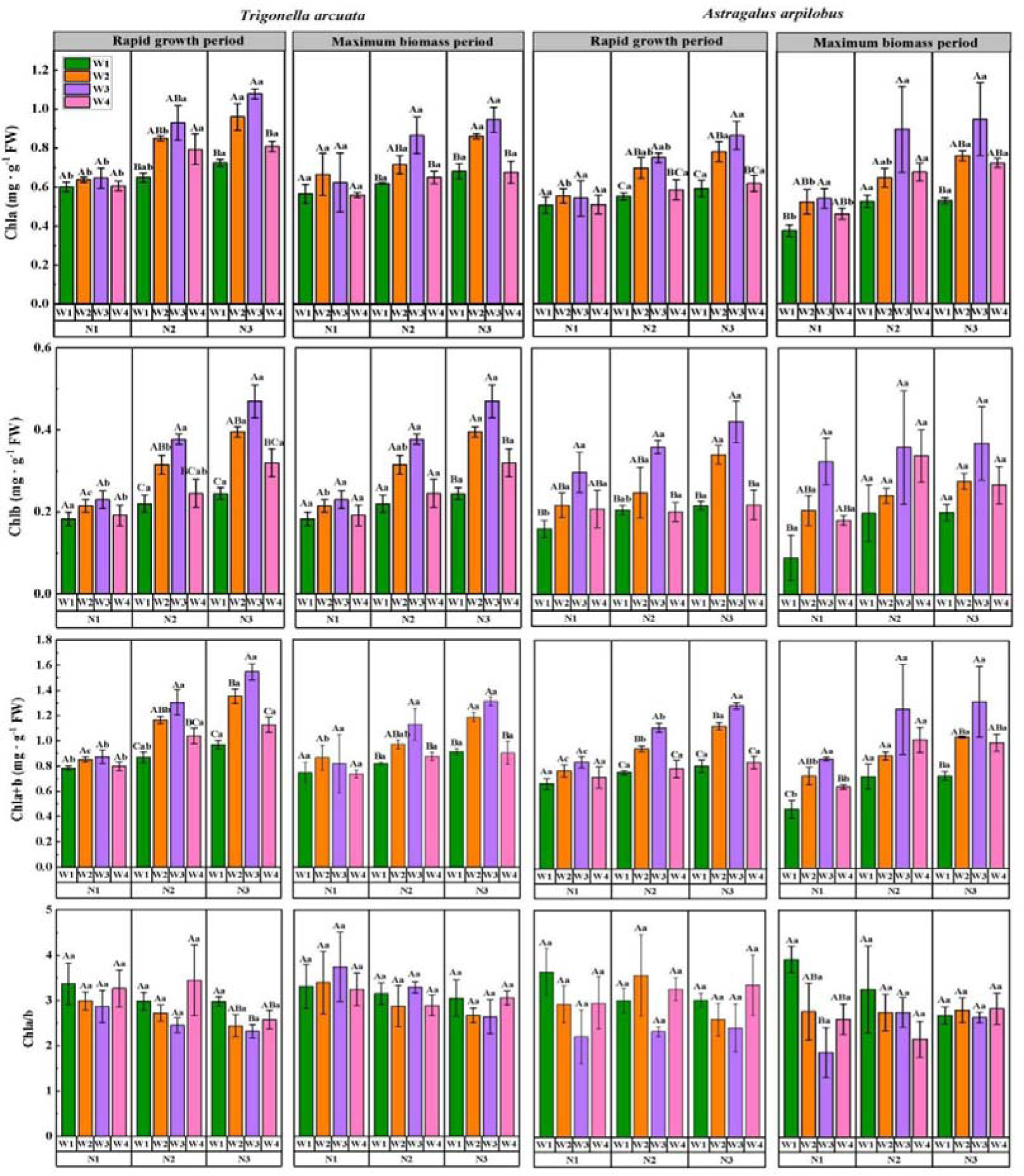
Chlorophyll content of two leguminous plants under the interaction of moisture and nitrogen. Different uppercase letters signify significant differences in the same plant under the same nitrogen level across various moisture gradients, and different lowercase letters indicate significant differences in the same plant under the same moisture gradient across different nitrogen gradients.

During different growth seasons, the Fv/Fm and YII of both plants were also significantly affected by moisture and nitrogen(Table 1). Under the same nitrogen addition, Fv/Fm and YII of both plants were mostly influenced by different moisture gradients, typically showing the trend: W3 > W2 > W4 > W1, and under the same moisture treatment, Fv/Fm and YII were influenced by different nitrogen levels, generally showing: N1 < N2 < N3. Combining the effects of moisture and nitrogen interaction, it was observed that Fv/Fm and YII of both leguminous plants were highest in the W3N3 treatment during both growth seasons, and lowest in the W1N1 treatment, with both indices being greater during the rapid growth period than at the maximum biomass period. In the more sensitive rapid growth period, Fv/Fm and YII of *Trigonella arcuata* in the W3N3 treatment were 1.40 and 1.57 times that of the W1N1 treatment, respectively, while for *Astragalus arpilobus*, they were 1.38 and 1.41 times, respectively. The patterns of variation in response to moisture and nitrogen were similar to those observed for chlorophyll content(Fig. 4).

**Fig. 4.**
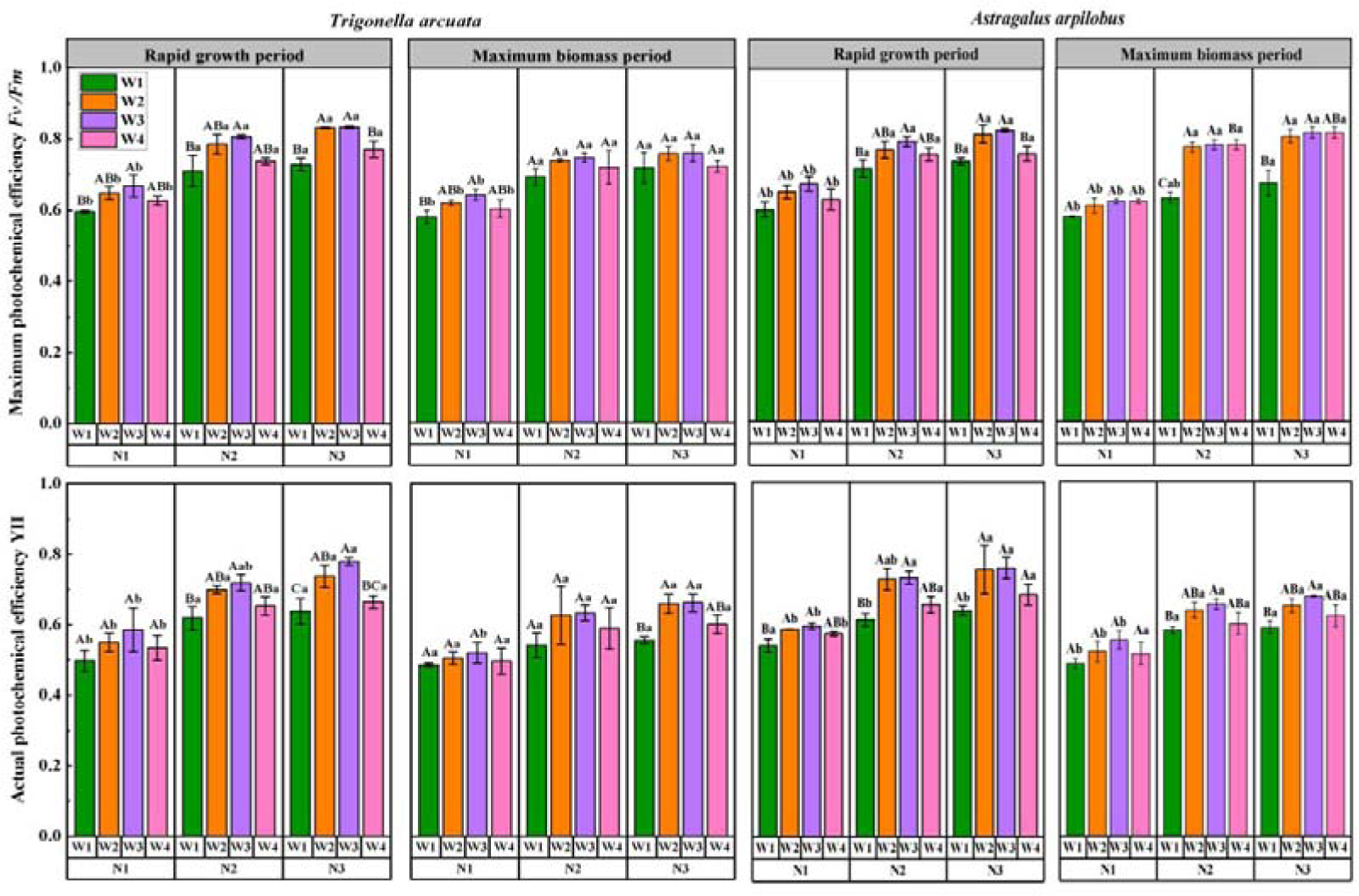
Chlorophyll fluorescence parameters of two leguminous plants under the interaction of moisture and nitrogen. Different uppercase letters represent significant differences in the same plant under the same nitrogen level across various moisture gradients, while different lowercase letters represent significant differences in the same plant under the same moisture gradient across different nitrogen gradients.

The results indicate that increasing gradients of moisture and nitrogen have a pronounced enhancing effect on the Chla, Chlb, Chla+b, Fv/Fm, and YII values of the two desert leguminous plants, with the interaction between moisture and nitrogen having a more prominent impact. Among these, the photosynthetic system of *Trigonella arcuata* was more sensitive to moisture and nitrogen, especially during the rapid growth period.

Osmotic regulation is an important physiological mechanism for plants to adapt to adverse stress conditions. Plants can actively accumulate various organic and inorganic substances to increase the concentration of cell sap, reduce osmotic potential, and enhance cell moisture retention, thereby improving plant adaptability to adverse conditions. In this study, species, growth season, moisture, nitrogen, species × moisture, growth season × moisture, growth season × nitrogen, and moisture × nitrogen significantly affected the free proline(Pro) content in the two plants(*P* < 0.001, *P* < 0.01, *P* < 0.05). Species, growth season, moisture, species × growth season, species × moisture, species × nitrogen, growth season × moisture, moisture × nitrogen, species × growth season × moisture, species × growth season × nitrogen, and species × moisture × nitrogen significantly influenced the soluble sugar(SS) content of the two plants(*P* < 0.001, *P* < 0.01, *P* < 0.05). The soluble protein(SP) content was more sensitive to these factors, with species, growth season, moisture, nitrogen, species × growth season, species × moisture, species × nitrogen, growth season × moisture, growth season × nitrogen, and moisture × nitrogen all having a significant impact on the SP content of the two plants(*P* < 0.001, *P* < 0.01, *P* < 0.05)(Table 1). Further analysis of the key factors, moisture and nitrogen, on the osmoregulation system of the two leguminous plants revealed that under the same nitrogen treatment, the Pro and SP contents of both plants were influenced by the moisture gradient. The Pro content was highest in the W1 treatment, followed by W3 and W2 treatments, and lowest in the W4 treatment. The SP content was generally higher in the W3 and W2 treatments and lower in the W4 and W1 treatments. Under the same moisture treatment, the Pro content of both plants under different nitrogen levels showed, in the W1, W2, and W3 treatments, a trend of N1 > N3 and N2, and in the W4 treatment, N1 < N2 < N3. The SS content did not follow a consistent pattern under the W1 and W2 treatments, but showed an opposite trend under the W3 and W4 treatments, with N1 > N2 > N3 in W3 treatment and N1 < N2 < N3 in W4 treatment, similar to the Pro content. The SP content, influenced by nitrogen levels, exhibited a more consistent pattern, generally showing: N1 < N2 < N3. Combining the effects of moisture and nitrogen interaction, it was found that the Pro and SS contents of both leguminous plants during the two growth seasons were highest in the W1N1, W1N3, and W1N2 treatments, and lowest in the W4N1 treatment. The SP content was highest in the W3N3 treatment and lowest in the W4N1 treatment, generally showing: rapid growth period < maximum biomass period. Notably, during the more sensitive maximum biomass period, the maximum Pro content of *Trigonella arcuata* and *Astragalus arpilobus* in the W1N3 treatment was 4.88 and 3.45 times the minimum value in the W4N1 treatment, respectively. The SS content of *Astragalus arpilobus* in the W1N3 treatment was 2.70 times that of the W4N1 treatment. The SP content of both plants in the W3N3 treatment was 2.37 and 2.51 times that of the W4N1 treatment, respectively(Fig. 5).

**Fig. 5.**
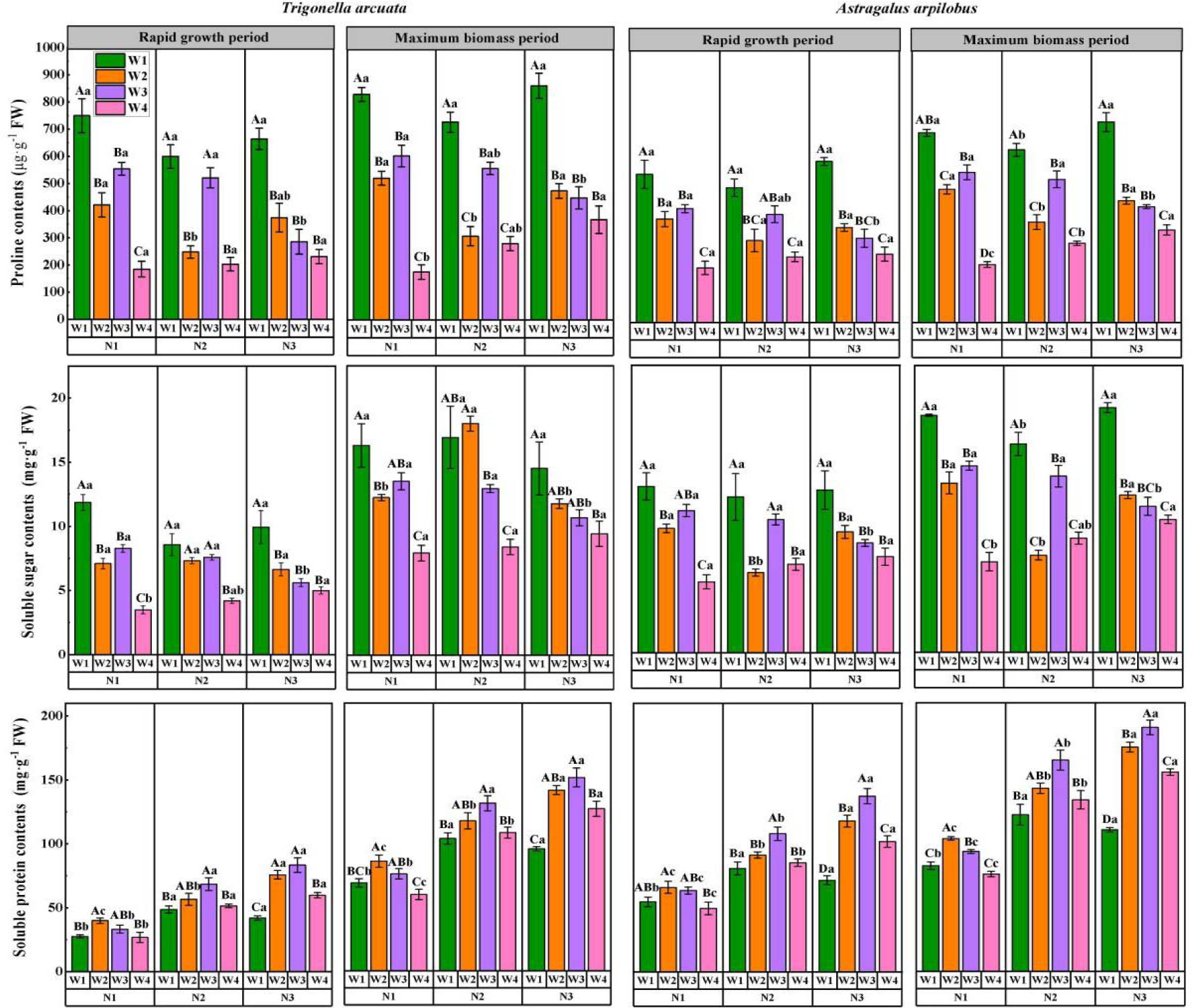
Osmoregulatory substance content of two leguminous plants under the interaction of moisture and nitrogen. Different uppercase letters indicate significant differences in the same plant under the same nitrogen level across different moisture gradients, and different lowercase letters signify significant differences in the same plant under the same moisture gradient across different nitrogen gradients.

The results indicate that the Pro and SS contents of the two plants are influenced by the interaction of moisture and nitrogen in a similar pattern. Under drought and low nitrogen conditions, desert leguminous plants increase their Pro and SS contents to cope with adverse stress. In contrast, the changes in SP are opposite; the increase in moisture and nitrogen accelerates the accumulation of SP content in the plants. A comprehensive analysis found that the sensitivity of the osmoregulatory substances in the two leguminous plants to moisture is in the order of Pro > SS > SP, while their sensitivity to nitrogen is in the order of SP > Pro > SS. This indicates that the osmoregulatory mechanisms in these plants are differentially influenced by environmental factors, with Pro being the most sensitive to moisture stress.

Drought and nitrogen deposition can lead to the generation of a large amount of Reactive Oxygen Species(ROS) in plant cells, causing increased lipid peroxidation and severely damaging the composition and integrity of the cytoplasmic membrane. This is primarily indicated by the accumulation of malondialdehyde(MDA), a lipid peroxidation product. In addition to osmotic regulation, plants activate an antioxidant enzyme defense system to scavenge excess reactive oxygen species, thereby combating stress conditions. The Superoxide Dismutase(SOD), Peroxidase(POD), and Catalase(CAT) are particularly important protective enzymes in this antioxidant system.

In this study, the growth season, moisture, nitrogen, and moisture × nitrogen significantly influenced the MDA in both plant species(*P* < 0.001). The species, growth season, moisture, nitrogen, and moisture × nitrogen had a highly significant effect on SOD content(*P* < 0.001). The POD content was more sensitive to these factors, with species, growth season, moisture, nitrogen, species × growth season, species × moisture, growth season × moisture, moisture × nitrogen, species × moisture × nitrogen, growth season × moisture × nitrogen, and species × growth season × moisture × nitrogen all showing a highly significant influence on POD content in both plants(*P* < 0.001). Additionally, the species, growth season, moisture, nitrogen, species × moisture, growth season × moisture, moisture × nitrogen, species × growth season × moisture, growth season × moisture × nitrogen, and species × growth season × moisture × nitrogen significantly affected CAT content(*P* < 0.001, *P* < 0.01, *P* < 0.05)(Table 1). Further analysis of the key factors, moisture, and nitrogen, on the cell membrane system and antioxidant enzyme system of the two leguminous plants revealed that under the same nitrogen addition, the MDA, SOD, POD, and CAT contents of both plants were mostly influenced by moisture gradients across three nitrogen levels, typically showing: W1 > W3 > W2 > W4, with only MDA and CAT content at the N3 level showing W1 > W2 > W3 > W4. Under the same moisture treatment, the MDA and SOD contents influenced by nitrogen level showed: N1, N3 > N2 in W1 and W2 treatments, N1 > N2 > N3 in W3 treatment, and the opposite trend in W4 treatment: N1 < N2 < N3. The POD content, influenced by moisture gradient, showed a similar pattern in W1, W2, and W3 treatments: N1 > N3, N2, while the variation in W4 treatment was consistent with MDA and SOD. CAT content, influenced by moisture gradient, exhibited a similar trend in W1 and W4 treatments: N3 > N1 > N2, and a similar pattern in W2 and W3 treatments: N1 > N3, N2. Combining the effects of moisture and nitrogen interaction, it was observed that in most cases, the MDA, SOD, POD, and CAT contents of both plants were highest in the W1N3 and W1N1 treatments, and lowest in the W4N1 and W2N2 treatments. Only MDA content showed a pattern controlled by moisture and nitrogen across the two growth seasons: rapid growth period < maximum biomass period, while SOD, POD, and CAT showed the opposite pattern: rapid growth period > maximum biomass period(Fig. 6).

**Fig. 6.**
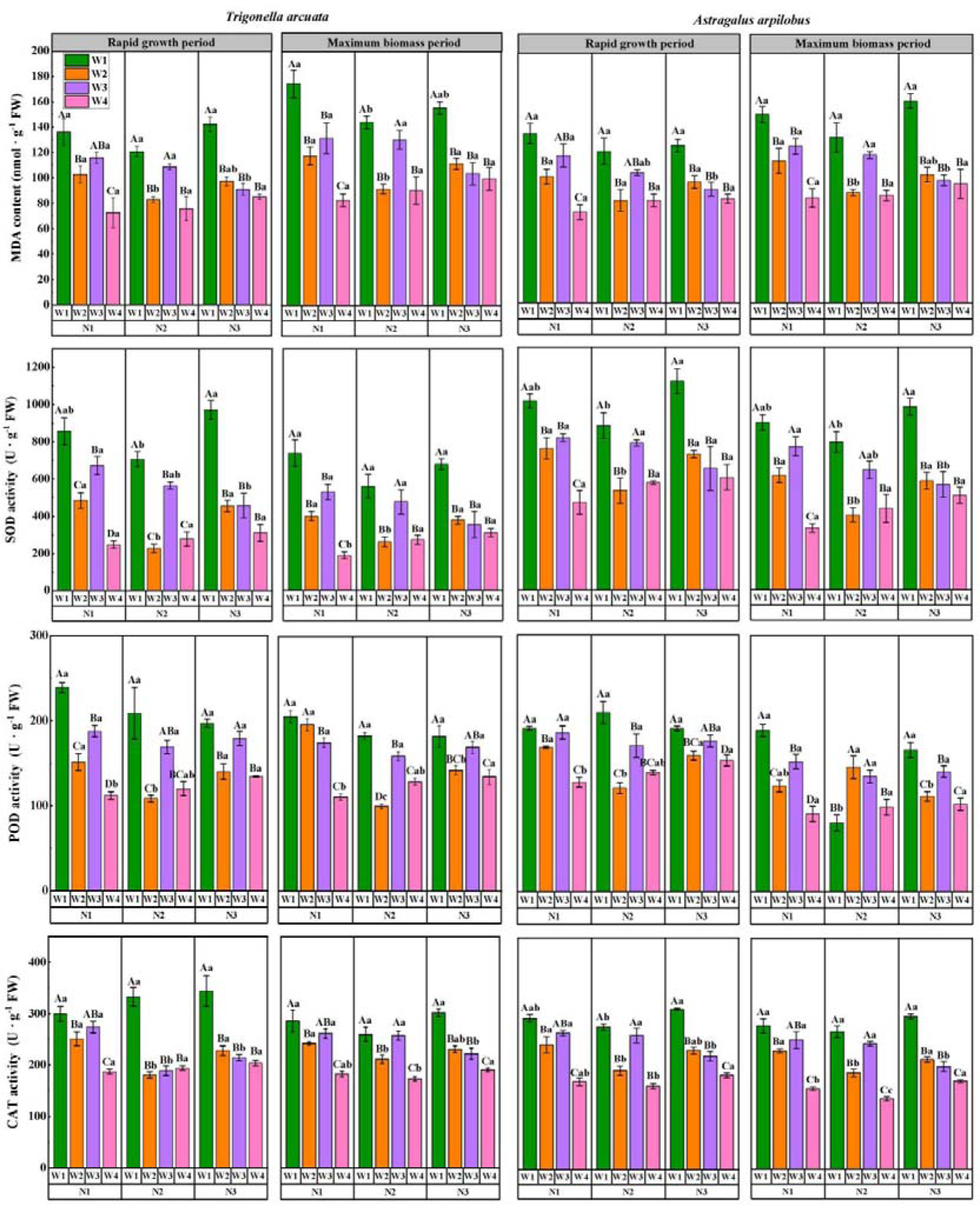
Antioxidant enzyme content of two leguminous plants under the interaction of moisture and nitrogen. Different uppercase letters indicate significant differences in the same plant under the same nitrogen level across different moisture gradients, while different lowercase letters signify significant differences in the same plant under the same moisture gradient across different nitrogen gradients.

**Fig. 7.**
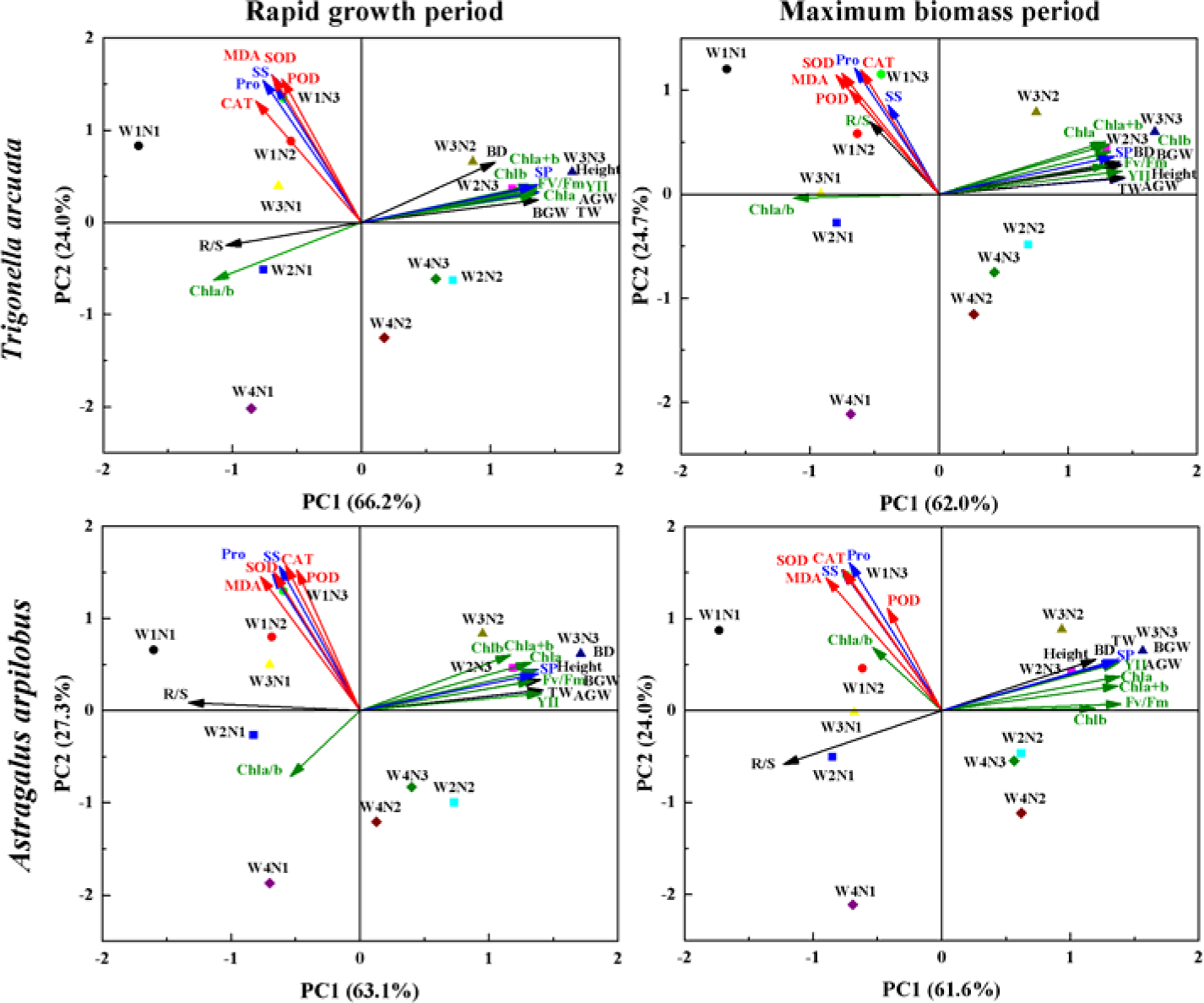
Principal component analysis of correlations between growth characteristics, chlorophyll content, osmoregulatory substance content and indicators of antioxidant enzyme system of 2 plants under different moisture and nitrogen treatments. The black arrows represent the growth characteristics, the green arrows represent the chlorophyll content characteristics, the blue arrows represent the osmoregulatory substance content characteristics, and the red arrows represent the indicators of cell membrane and antioxidant enzyme system.

**Fig. 8.**
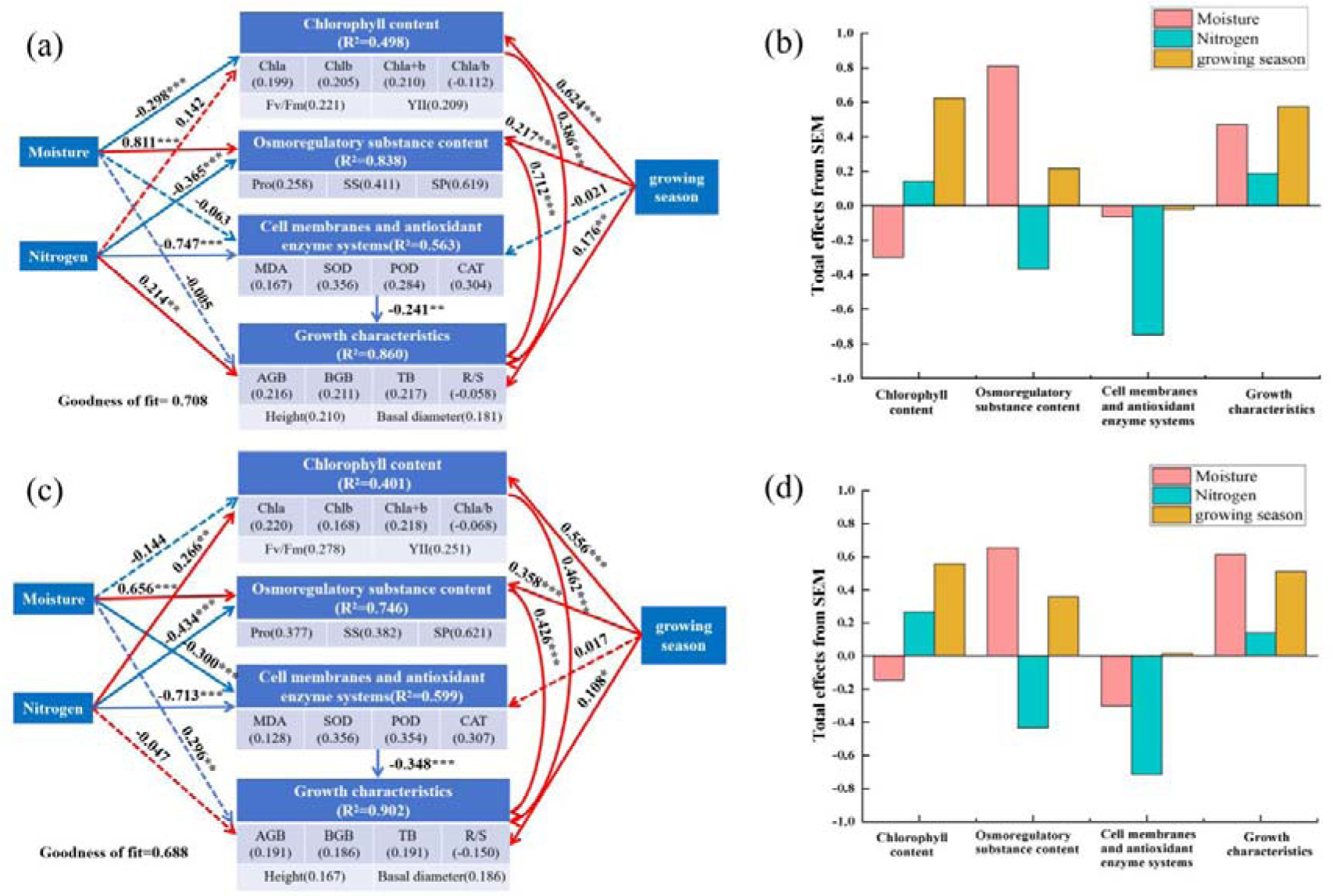
PLS-PM pathway analysis of the effects of moisture, nitrogen and growing season on growth characteristics, chlorophyll content, osmoregulatory substance content and antioxidant enzyme systems of two plant species. Where a and b are effect relationship plots for Trigonella arcuata and c and d are effect relationship plots for Astragalus arpilobus. Arrows indicate causal relationships:red lines indicate positive effects and blue lines indicate negative effects. Numbers next to arrows represent standardised path coefficients. Significance levels are indicated as:\**p* < 0.05, \*\**p* < 0.01, \*\*\**p* < 0.001.

The results indicate that drought and low nitrogen stress also increase the accumulation of protective enzymes in the cell membrane system and antioxidant enzyme system to cope with membrane damage and reactive oxygen species accumulation. During the rapid growth period, the cell membrane defense system is not as robust as during the maximum biomass period. However, plants can eliminate ROS through the accumulation of the antioxidant enzyme defense system to combat stress conditions, exhibiting a phenomenon of moisture diluting the nitrogen supply effect. Please continue the translation following the same standards.

### PCA and path analysis

PCA analysis was used to determine the correlations and key influencing factors between the indicators of growth characteristics, chlorophyll content, osmoregulatory substance content and antioxidant enzyme system of the two plants. Sorting analysis explained 92%, 86.7%, 90.4%, and 85.6% of the total variance distribution, respectively. The combined analysis revealed positive correlations between above-ground, below-ground, and total fresh weight, plant height, basal diameter, and Chla, Chlb, Chla+b, Fv/Fm, YII, and SP of the two species, while they were negatively correlated with R/S and Chla/b, both in the maximum biomass and maximum biomass periods. Indicators of cell membrane and antioxidant enzyme system(MDA, SOD, POD, CAT) were positively correlated with Pro and SS, while they were not strongly correlated with other indicators. Comprehensive analysis of the correlations between treatment groups and indicators under the influence of moisture and nitrogen interactions revealed that aboveground, belowground and total fresh weight, plant height, basal diameter and Chla, Chlb, Chla+b, Fv/Fm, YII, and SP were positively correlated with W2N2, W3N2, W3N3, W4N2, W4N3, with the strongest correlations with W3N3 and W2N3. while the cell membrane and antioxidant enzyme system indicators(MDA, SOD, POD, CAT), Pro and SS were positively correlated with W1N1, W1N2, W1N3, W2N1, W3N1, W4N1, with the strongest correlations with W1N3 and W1N2.

In order to minimise confounding interactions between causal factors, PLS-PM was used to further determine the direct and indirect effects between moisture, nitrogen and growing season on growth characteristics, chlorophyll content, osmoregulatory substance content and antioxidant enzyme systems of the two plant species. The goodness of fit indicated that the mean predictive values of the model were 0.708 and 0.688. Moisture had a negative effect on chlorophyll content(−0.298), cell membrane and antioxidant enzyme system(−0.063) and a significant positive effect on osmoregulatory substances(0.811) and growth traits(0.472) of *Trigonella arcuata*; Nitrogen had a positive effect on chlorophyll content(0.142) and growth characteristics(0.188) and a negative effect on osmoregulatory substances(−0.365) and on cell membranes and antioxidant enzyme systems(−0.747) in *Trigonella arcuata*; the growing season had a significant positive effect on chlorophyll content(0.624), osmoregulatory substances(0.811) and growth characteristics(0.472) in *Trigonella arcuata*. Osmoregulatory regulating substances(0.217) and growth characteristics(0.577) positively, while the effect on cell membrane and antioxidant enzyme systems was not significant.

Analysing the direct and indirect effects of moisture, nitrogen and growing season on growth characteristics, chlorophyll content, osmoregulatory substances content and antioxidant enzyme system of *Astragalus arpilobus*, it was found that moisture had a negative effect on chlorophyll content(−0.144), cell membrane and antioxidant enzyme system(−0.300) and a negative effect on osmoregulatory substances(0.656) and growth characteristics(0.613); nitrogen had a positive effect on chlorophyll content(0.266) and growth characteristics(0.139) of *Astragalus arpilobus*, while it had a negative effect on osmoregulatory substances(−0.434) and on cell membrane and antioxidant enzyme systems(−0.713); Growing season positively affected chlorophyll content(0.556), osmoregulatory substances(0.358) and growth characteristics(0.512) of *Astragalus arpilobus*, whereas the effect on cell membrane and antioxidant enzyme systems was not significant. Overall, increase in water positively affected osmoregulatory substances content and growth characteristics of the two plants, while negatively affecting chlorophyll content and cell membrane and antioxidant enzyme systems. Increase in nitrogen level positively affected chlorophyll content and growth characteristics, while negatively affecting osmoregulatory substances content and cell membrane and antioxidant enzyme systems. Whereas, accumulation of growing season had positive effect on chlorophyll content, growth characteristics and osmoregulatory substances content.

## Discuss

Legumes contribute absolutely to the nitrogen input to terrestrial ecosystems due to their typical rhizomatous nitrogen fixation and are effective nitrogen input centres for nitrogen-poor desert ecosystems(Peoples et al., 2015; Rasmussen et al., 2019; Zhang et al., 2018). However, the alteration of precipitation patterns and the increase of nitrogen deposition under the influence of global climate change have profoundly affected the production and functioning of desert ecosystems(Huang et al., 2016). Based on the importance of legumes in desert ecosystems and the threat of altered precipitation and increased nitrogen deposition faced by desert ecosystems, we chose ephemeral legumes, which are extremely sensitive to responding to environmental changes for our study, and posed the scientific question: how do the growth and physiological functions of ephemeral legumes with nitrogen-fixing functions respond to the changes in moisture and nitrogen conditions in a typical temperate desert ecosystem? We attempted to trace back from the phenological characteristics to the physiological and biochemical levels to comprehensively analyse the coupled effects of moisture and nitrogen on the growth of leguminous plants.

### Effects of moisture-nitrogen coupling on growth accumulation of ephemeral legumes

Moisture and nitrogen, as the two most important factors limiting plant growth in the Gurbantunggut Desert(Hooper and Johnson, 1999), and the climatic context of warming and humidification and increased nitrogen deposition make it difficult to clarify the direction of development of ecosystem stability in the region. Moisture is an important component in maintaining the morphology of the plant body, and a transporter and participant in a series of physiological and biochemical activities within the plant(Chaves et al., 2003; Flexas et al., 2004; Lawlor, 2002; Zhu, 2002). Drought and moisture scarcity can limit normal plant growth, but excessive moisture supply can inhibit respiration in the plant root system and thus affect plant growth(Colmer and Voesenek, 2009; Pandey et al., 2000). Nitrogen is a vital element for plant growth, which can directly affect the morphology and physiological functions of organs such as roots, stems and leaves, and ultimately affect the accumulation of biomass(Liu and Von Wirén, 2017; Xu et al., 2012). Most of the studies have pointed out that appropriate nitrogen can promote the accumulation of photosynthetic pigments and increase the photosynthetic rate in plants, which is conducive to the storage of nutrients, while the lack of nitrogen and excessive nitrogen will inhibit the accumulation of related products to varying degrees, limiting the growth of plants(Hachiya and Sakakibara, 2017). Plants in arid zones usually evolve a variety of adaptive strategies to cope with unfavourable conditions such as drought and soil nutrient depletion. For example, under moisture deficit conditions, plants can increase the area and volume of the root system to access soil moisture by increasing their root-shoot ratio, and they can also adjust the area and thickness of the leaves, adjust the stomatal movement, and increase the epidermal hairs and the epidermal wax paper in order to reduce the evaporation of moisture from the leaf blades(Chaves et al., 2009; Comas et al., 2013; Farooq et al., 2009). In this study, aboveground, belowground and total fresh weights, plant height and basal diameter of *Trigonella arcuata* and *Astragalus arpilobus* were all affected by different moisture gradients under the same nitrogen addition treatments as W3 > W2, W4 > W1, and we found that excessive moisture (W4) also limited the growth of drought-tolerant plants, but not to the same extent as drought conditions (W1), while the root-shoot ratio of *Trigonella arcuata* and *Astragalus arpilobus* were mostly shown by the different moisture gradients. Most of the effects of moisture gradient were shown as W1 > W4 > W3 and W2, indicating that drought and excess moisture would directly affect the direction of nutrient flow in both above-ground and below-ground parts of the plant. The response of plants to exogenous nitrogen is also very sensitive, and the increase of nitrogen also accelerates soil mineralisation, promotes microbial metabolism, facilitates seed germination and seedling growth, and promotes rapid renewal and reproduction of herbaceous plant communities(Galloway et al., 2008; LeBauer and Treseder, 2008; Vitousek et al., 2009). However, excess nitrogen can also alter the photosynthetic rate, chlorophyll content, cell membrane permeability, intracellular pH and nutrient balance and other metabolic processes. Since nitrogen tolerance varies among species, exogenous nitrogen inputs can affect competition and coexistence among plants, further affecting the structural stability of ecosystems(Bobbink et al., 2010; De Schrijver et al., 2011; Stevens et al., 2011). In this study, the aboveground, belowground and total fresh weight, plant height and basal diameter of the two plants were affected by different nitrogen levels under the same moisture level: N1 < N2 < N3, indicating that the increase of nitrogen level directly promoted the nutrient accumulation of plants. The root-shoot ratio was regulated by nitrogen levels in contrast to that in low nitrogen environments, plants would accumulate more energy in the roots. We comprehensively analysed the indicators under moisture-nitrogen coupling and found that the coupling of moisture and nitrogen had both beneficial and detrimental effects on plant growth, with above-ground, below-ground and total fresh weights, plant heights and basal diameters of the two plants being smaller in the W4N1 and W1N1 treatments, and largest in the W3N3 treatment. As we expected, sufficient moisture accelerated the nutrient metabolism process in the plants and promoted morphogenesis and biomass accumulation. In addition each index was greater than W4N3, W4N2, and W4N1 at W3N3, W3N2, and W3N1 treatments, respectively, which is similar to the conclusion of most studies that excess moisture hinders nitrogen uptake by plants. We hypothesised that when the soil moisture content was too high, it would inhibit the respiration of the root system, thus affecting its access to soil nitrogen sources. However, all indicators were greater in W4N3 treatment than in W4N2 treatment, and we found that when faced with nitrogen excess and nitrogen accumulation, sufficient moisture has a diluting effect on both nitrogen uptake by the plant and nitrogen content in its own body, so that oxidative stress caused by excess nitrogen accumulation can be avoided. However, at the N1 level, the plants were in nitrogen starvation, at which time the effect of moisture factor on the plants was very small, both of which were unfavourable to the morphogenesis and energy accumulation of the plants. Each index was significantly smaller than W4N3, W2N3, and W3N3 in W1N3 treatment, and we believe that drought also hinders nitrogen uptake and transport in turn.

### Physiological processes and key drivers of ephemeral legumes in response to moisture and nitrogen stresses

In addition to analysing the phenological and morphological characteristics of potted legumes under the interactive effects of moisture and nitrogen, we further combined physiological and biochemical means to explore the characteristics of the photosynthetic system, osmoregulatory system, cell membrane system, and antioxidant enzyme system of potted legumes in response to the changes of moisture and nitrogen at a deeper level. Chlorophyll is an important pigment for photosynthesis in plants, and its content not only represents the intensity of photosynthesis, but also reflects the growth and development of plants under adversity stress(Kalaji et al., 2017; Murchie and Lawson, 2013). Studies have shown that drought stress causes stomatal closure, affecting transpiration and limiting CO_2_ uptake, while at the same time causing heat stress in leaves, which in turn affects photosynthetic output, resulting in a reduction in chlorophyll content and a slowing down of photosynthesis rate(Lawlor and Tezara, 2009). Nitrogen, as a key element of chlorophyll molecule, and insufficient nitrogen supply will directly affect chlorophyll synthesis and reduce photosynthetic efficiency(Paul and Foyer, 2010). In this study, the increase of moisture and nitrogen gradient had a significant effect on the Chla, Chlb, Chla+b, Fv/Fm, and YII values of the two desert legumes, and the effect of moisture-nitrogen interaction was more prominent. However, the chlorophyll content, photosynthesis efficiency and electron transfer capacity were significantly lower under W4 treatment compared with W2 and W3, and we speculated that the high soil moisture content made the roots insufficiently oxygenated, thus affecting the nutrient uptake capacity of the plants, which indirectly led to the decrease of chlorophyll content and photosynthesis efficiency. The Chla/b ratio increased when moisture was reduced, probably because Chla degradation was slower than Chlb. Osmoregulation is an important physiological mechanism for plant adaptation to adversity stress, and the plant body can increase the concentration of the cytosol, reduce the osmotic potential, and improve the cellular moisture-holding capacity through the active accumulation of a wide range of organic and inorganic substances, thus enhancing the plant resistance to adapt to adversity(Chaves et al., 2009; Yamaguchi and Blumwald, 2005). It has been pointed out that when drought restricts the growth of plants, in order to avoid tissue damage, plants will synthesise a large amount of proline and a small amount of soluble sugar to regulate the osmotic balance of the cell, and nitrogen as a key element in protein synthesis, the addition of exogenous nitrogen has a promotional effect on the synthesis of soluble proteins, but the high nitrogen will damage the cellular structure, affecting the synthesis of proteins, which will have an inhibitory effect on the growth of the plant(Ashraf and Foolad, 2007). In this study, during drought and low nitrogen treatments, desert legumes would respond to the adversity stress by increasing their Pro and SS contents, while the SP changes were the opposite, except for the W4 treatment, the increase of moisture and nitrogen accelerated the accumulation of SP content in the plants, even under high nitrogen, which, in our opinion, may be related to the inherent nitrogen scarcity in this study area and the strong nitrogen utilisation ability of legumes closely related. In addition to osmoregulation, plants can also produce a large amount of ROS when they are subjected to drought and other adversity stresses, causing damage to the plant membrane system. In order to prevent cellular damage, the increase of protective enzymes, such as MDA, CAT, POD, and SOD in the plant cells promotes to increase the accumulation of the corresponding protective enzymes in the cell membrane system and the antioxidant enzyme system in order to cope with the membrane damage and the accumulation of reactive oxygen species(Apel and Hirt, 2004; Gill and Tuteja, 2010; Miller et al., 2010). In this study, the MDA, SOD, POD, and CAT contents of the two plants possessed the maximum values in W1N3 and W1N1 treatments, and the minimum values in W4N1 and W2N2 treatments. Drought and low nitrogen stress would increase the contents of MDA, CAT, POD, and SOD in order to balance the ROS, but the lack of oxygen in the roots during the W4 treatment led to the slowing down of the cellular metabolic activities, and the production of ROS might be be reduced, thus decreasing the demand for antioxidant enzymes such as CAT, POD, and SOD.

PLS-PM pathway analysis showed that the negative effects of chlorophyll content, cell membrane and and antioxidant enzyme systems in response to excess moisture(W4) were very obvious in the two plants. Although Pro and SS among the osmoregulatory substances were also negatively correlated with the increase in moisture, the greater contribution of the positive effect of SP led to an overall positive correlation between the increase in moisture and osmoregulatory substance content, which in turn indirectly positively affected the growth characteristics of the plants. The osmoregulators in ephemeral legumes were the most sensitive to moisture than the other indicators, which may be closely related to the fact that osmoregulators can directly regulate the intracellular moisture status and osmotic pressure, and the process of regulating them is more rapid and efficient in the event of moisture stress. Nitrogen increase was negatively correlated with osmoregulators, cell membranes and antioxidant enzymes, which better confirms that under low nitrogen conditions, plants will improve their related physiological indicators of stress to cope with unfavourable environments(Kusano et al., 2011), and that excess nitrogen has a lesser effect on ephemeral legumes than low nitrogen environments, and that cell membranes and antioxidant enzymes respond more significantly to nitrogen stress in ephemeral legumes than in other systems. We hypothesise that too little and too much nitrogen directly affects nitrogen metabolism, oxidative stress and resource reallocation processes in plants, and that the rapid response of the plant’s cell membranes and antioxidant enzyme systems is a direct mechanism for regulating the above processes(Foyer and Noctor, 2003; Stitt and Krapp, 1999). Overall, the increase in moisture and nitrogen levels indirectly affected the apparent growth characteristics of the two plants through the regulated photosynthetic chlorophyll system, osmoregulatory system, cell membrane and antioxidant enzyme systems, among which the growth traits(above-ground, below-ground and total fresh weights, plant height, basal diameter) and chlorophyll content indicators (Chla, Chlb, Chla+b, Fv/Fm, YII), as well as the SP content in response to moderate increase in moisture and nitrogen(W3N3 and W2N3) were positive, whereas cell membrane and antioxidant enzyme system indicators(MDA, SOD, POD and CAT), and osmotic regulators (Pro and SS) were susceptible to accumulating during exposure to extreme moisture and nitrogen stresses to enhance cellular resistance.

### Growing season differences in physiological and metabolic processes in ephemeral legumes

In plant growth and development, environmental differences due to seasonal changes directly affect aspects of photosynthetic efficiency, phytohormone levels, metabolic activities, and adversity response mechanisms, which in turn influence plant growth traits(Chaves et al., 2009). Plants also adapt to seasonal changes by adjusting their metabolic activities, including photosynthesis, respiration, and the synthesis of secondary metabolites, and these adjustments reflect altered energy balance and growth dynamics in plants(Cleland et al., 2007; Estiarte and Peñuelas, 2015; Morin et al., 2010; Körner and Basler, 2010). In this study, we attempted to analyse the changes in the regulation of physiological and metabolic activities in the plant itself in response to stress at different stages of growth and development by excluding the effects of external environmental factors. We compared each index of *Trigonella arcuata* and *Astragalus arpilobus* during the rapid growth period and the maximum biomass period and found that the chlorophyll content indexes (Chla, Chlb, Chla+b, Fv/Fm, and YII), and antioxidant enzyme system indexes (SOD, POD, and CAT) of the two plants behaved in the following ways: rapid growth period > maximum biomass period. We suggest that during the rapid growth period, seedlings are at a critical stage of morphogenesis, and plants are actively allocating resources to support new leaf formation and expansion, as well as other growth activities. Higher chlorophyll content contributes to enhanced photosynthesis and supports rapid energy and biomass accumulation(Porcar-Castel et al., 2014). Meanwhile, high values of Fv/Fm and YII as indicators of photosynthetic efficiency during the rapid growth period indicate the efficient energy conversion and electron transfer capacity of photosynthesis system II, whereas by the period of maximum biomass, plant growth rate slows down, and photosynthesis and resource allocation may be shifted more towards supporting the maintenance of existing tissues and reproductive growth(Baker, 2008; Maxwell and Johnson, 2001). As a result, chlorophyll content and photosynthetic efficiency metrics may decline relative to the rapid growth period. The cellular metabolic activities of plants are significantly enhanced during the rapid growth phase, resulting in a high production of ROS, and plants enhance their oxidative stress defences by continuously increasing the enzyme activities of their own antioxidant systems. By the later stages of growth, plants have established more stable defence mechanisms, thus reducing the need for immediate response to oxidative stress(Cleland et al., 2007; Estiarte and Peñuelas, 2015; Körner and Basler, 2010; Morin et al., 2010). In contrast, growth traits(aboveground, belowground, and total fresh weight, plant height, basal diameter) and osmoregulatory substances (Pro, SS and SP) and MDA content of cell membrane systems were shown to be the following: rapid growth period < maximum biomass period. We hypothesise that when plants develop to the period of maximum biomass, the development of relevant growth indicators slows down and has reached a stable maturity level, when biomass accumulation peaks and the physiological senescence process begins, which affects the metabolic activities and growth potential of the cells. Cell membrane integrity may be reduced by senescence-related oxidative damage, thus leading to increased MDA content. As the root system ages and soil nutrients become progressively depleted, plants may experience greater moisture and nutrient stress, which requires plants to adapt their physiological mechanisms to utilise limited resources more efficiently, including improving cellular osmotic balance and preserving cellular structures by increasing the synthesis of osmoregulatory substances(Serraj and Sinclair, 2002; Verslues et al., 2006). According to PLS-PM pathway analysis, it is known that chlorophyll content indicators and growth characteristic indicators are most sensitive in response to changes in the growing season, and that the differences between plants in different growing seasons reflect changes in the intensity of their photosynthesis and energy requirements. The adaptive characteristics of physiological and metabolic processes in plants to seasonal changes can more comprehensively explain the process of environmental factors on plant growth.

## Conclusions

The results showed that moisture-nitrogen coupling explained the physiological processes of ephemeral legumes better than single-factor effects, and drought also impeded nitrogen uptake and transport, whereas sufficient moisture accelerated the nutrient metabolic processes of plants, and there was nutrient dilution once moisture was in excess, but the moisture factor had little effect on plants under nitrogen starvation. The osmoregulatory substances content of the two plants was the most sensitive to moisture, while the cell membrane and antioxidant enzyme systems were the most sensitive to nitrogen stress. Chlorophyll content and growth characteristics were the most sensitive to seasonal variations, and the differences between growing seasons reflected the changes in photosynthetic intensity and energy requirements. This study can provide a theoretical basis for understanding the response characteristics of desert legumes to nitrogen deposition and precipitation under global climate and environmental changes, and to nitrogen cycling in the region.

## Author contributions

Yuxin Xiao, Boyi Song, and Weiwei Zhuang: conceptualization;Yuxin Xiao and Boyi Song: writing the initial draft; Jinqiu Li, Nargiza Galip, Ao Yang and Xinyu Zhang: revising the manuscript. All the authors read and approved the final version of the manuscript.

